# Assessing the effectiveness of oxathiapiprolin towards *Phytophthora agathidicida*, the causal agent of kauri dieback disease

**DOI:** 10.1101/2021.03.10.434845

**Authors:** Randy F. Lacey, Michael J. Fairhurst, Kaitlyn J. Daley, Te Amohaere Ngata-Aerengamate, Haileigh R. Patterson, Wayne M. Patrick, Monica L. Gerth

**Author notes:** Corresponding author: M. L. Gerth.

## Abstract

*Phytophthora* species cause disease and devastation of plants in ecological and horticultural settings worldwide. A recently identified species, *P. agathidicida*, infects and ultimately kills the treasured kauri trees that are endemic to New Zealand. Currently there are few options for controlling or treating *P. agathidicida*. In this study, we sought to assess the toxicity of the oomycide oxathiapiprolin against several lifecycle stages of two geographically distinct *P. agathidicida* isolates. Half maximal effective concentration (EC_50_) values were determined to be approximately 0.1 ng/ml for inhibiting mycelial growth, indicating that *P. agathidicida* mycelia are more sensitive to oxathiapiprolin than those from most other *Phytophthora* species that have been studied. Oxathiapiprolin was also highly effective at inhibiting the germination of zoospores (EC_50_ = 2-9 ng/ml for the two isolates) and oospores (complete inhibition at 100 ng/ml). In addition, oxathiapiprolin delayed the onset of detached kauri leaf infection in a dose-dependent manner. Collectively, the results presented here highlight the significant potential of oxathiapiprolin as a tool to aid in the control of kauri dieback disease.

## Introduction

Members of the oomycete genus *Phytophthora* cause disease and destruction to plants in agricultural and natural ecosystems worldwide (Hansen et al. 2012). For example, it has been estimated that the infamous causal agent of potato late blight, *Phytophthora infestans*, continues to cause >$6 billion of damage yearly (Haverkort et al. 2008). In New Zealand, a recently identified species named *P. agathidicida* is threatening kauri (*Agathis australis*), which are treasured, long-lived native conifers (Bradshaw et al. 2020; Weir et al. 2015). Kauri are giant trees – the largest, Tāne Mahuta, has a trunk girth of over 13 m – and they have a vital ecological role as foundation species in the forests they inhabit (Wyse et al. 2014). *P. agathidicida* infects kauri of all ages *via* the roots, causing trunk lesions, canopy thinning, root and collar rot, and ultimately death (Bellgard et al. 2016). While kauri dieback disease was first documented in the 1970s, it has spread rapidly in the past decade and now poses a significant long-term threat to the species (Black et al. 2018; Waipara et al. 2013).

Currently, there are few options for controlling or treating *P. agathidicida*. The main tool for reducing pathogen spread is physical barriers such as walking track closures and shoe cleaning stations. The only chemical treatment that has been used in the field is phosphite (phosphorous acid). In trunk injection field trials, phosphite improved canopy health, and reduced lesion activity and expansion (Horner et al. 2015). However, potential phytotoxic effects (Bradshaw et al. 2020; Horner et al. 2015) and the potential for resistance to develop (Dobrowolski et al. 2008; Hao et al. 2020) are concerns regarding phosphite treatment. Other chemical controls have been explored (Lawrence et al. 2017), including surveying native New Zealand plants for their production of oomycides (Lawrence et al. 2019), but none of these options has been field tested to date.

Oxathiapiprolin is a first-in-class piperindinyl thiazole isoxazoline oomycide (Pasteris et al. 2016) that is highly effective against many oomycete species when applied preventatively or curatively (Belisle et al. 2019; Bittner et al. 2017; Cohen 2015; Cohen et al. 2018; Gray et al. 2018; Humann et al. 2019; Ji and Csinos 2015; Miao et al. 2016b; Miao et al. 2016a; Qu et al. 2016a). It can be applied *via* foliar sprays, soil applications (*e.g*. in-furrow) or soil drenches and it is efficiently translocated throughout plant tissues in a variety of horticultural species (Cohen 2015, 2020; Qu et al. 2016b), including perennial tree crops (Gray et al. 2020). In addition, oxathiapiprolin has low phytotoxic effects in host tissue. For example, in downy mildew infected sunflower, the highest tested concentrations of oxathiapiprolin were effective in treating disease while also causing no phytotoxic effects (Humann et al. 2019).

The intracellular target of oxathiapiprolin is an oxysterol binding protein (OSBP)-related protein (ORP) (Bittner et al. 2017; Miao et al. 2018; Miao et al. 2016b; Pasteris et al. 2016). While the precise function of ORPs in oomycetes is unknown, this family of proteins exists throughout eukaryotes and its members are involved in a broad range of functions including intracellular sterol transport, lipid metabolism, and signal transduction (Raychaudhuri and Prinz 2010).

In this study, we have annotated the sequence of the *P. agathidicida* ORP1 gene (*PaORP1*). In two geographically distinct isolates of *P. agathidicida*, we have assessed the efficacy of oxathiapiprolin against different stages in the lifecycle including mycelial growth, and both motility and germination of the infectious zoospores. By designing a protocol to purify the metabolically dormant survival spores (oospores), we were also able to test the efficacy of oxathiapiprolin against this life cycle stage, for the first time for any *Phytophthora* species. Overall, our work establishes a baseline that strongly supports field testing of oxathiapiprolin for the management of kauri dieback disease.

## Materials and Methods

### Materials

*P. agathidicida* isolates NZFS 3770 and NZFS 3772 were obtained from the culture collection held at Scion (Rotorua, New Zealand). *P. agathidicida* 3770 was originally isolated from Coromandel, New Zealand, while 3772 was isolated near Auckland, New Zealand (Studholme et al. 2016). Oxathiapiprolin (≥98% purity) was purchased from Carbosynth (Compton, Berskhire, UK). It was stored at 1 mg/ml in 100% dimethyl sulfoxide (DMSO) at −20 °C. To account for possible solvent effects, DMSO concentrations were standardized and DMSO-only controls were performed in all experiments. Pimaricin (2.5%, w/v, aqueous solution), rifampicin, pentachloronitrobenzene, β-sitosterol and fluorescein diacetate were from Sigma Chemical Co. (St. Louis, MO, USA). Ampicillin was from GoldBio (St. Louis, MO, USA). TOTO-3 iodide was from Invitrogen. Brightfield microscopy was performed using an Olympus CKX53 inverted light microscope with Olympus cellSens Standard software for image processing.

### Bioinformatics

The *ORP1* gene sequences from *P. capsici, P. infestans, P. ramorum* and *P. sojae* have been published in the patent literature (Andreassi et al. 2013). Each was used as the query sequence for a pairwise alignment with the *P. agathidicida* 3772 draft genome (Studholme et al. 2016) using Parasail 2.4.1 (Daily 2016), implemented within SnapGene 5.0.7. A candidate transcriptional start site and a likely intron were annotated using the *Phytophthora* consensus sequences described previously (Kamoun 2003). The translated sequence of the *Pa*ORP1 protein was analyzed using ProtParam (Gasteiger et al. 2005), and the domains were identified using the NCBI Batch Conserved Domain Search tool (Lu et al. 2020).

### Routine culture conditions

*P. agathidicida* was cultured at 22 °C in the dark, unless otherwise noted. *P. agathidicida* isolates were initially cultured on selection plates comprising cornmeal agar (BD Difco; 17 g/l) supplemented with pimaricin (0.001%, w/v), ampicillin (250 *μ*g/ml), rifampicin (10 *μ*g/ml) and pentachloronitrobenzene (100 *μ*g/ml). A 6 mm diameter section of the leading edge of mycelia was transferred to a clarified V8 agar plate.

Clarified V8 agar was prepared by mixing 200 ml of Campbell’s V8 juice with 2 g CaCO_3_ on a magnetic stirrer for 10 min at room temperature. Next, it was clarified by centrifugation at 7000*g* for 10 min. The clarified juice (200 ml) was added to 800 ml of distilled, deionized water (ddH_2_O) with 15 g of bacteriological grade agar (Formedium, Hunstanton, UK) and sterilized by autoclaving. The isolates were then maintained by routinely transferring them to new clarified V8 agar plates for the duration of the study.

### Mycelial growth inhibition testing

Mycelial growth inhibition was measured on potato dextrose agar plates (BD Difco; 39 g/l). Initially, 6 mm diameter agar plugs were taken from the leading edge of mycelial growth on clarified V8 plates and transferred to potato dextrose plates amended with varying concentrations of oxathiapiprolin. The plates were then incubated at 22 °C in the dark for five days, with images subsequently taken of each plate. For each replicate, six radial measurements were taken from the centre of the outermost point of growth and averaged. The concentration of oxathiapiprolin required to effect 50% growth inhibition (the EC_50_) was calculated using GraphPad Prism Version 8 by plotting log-transformed oxathiapiprolin concentration against the average radius of growth.

### Zoospore production and testing

To produce zoospores, ten 6 mm diameter agar plugs were taken from the leading edge of mycelial growth on clarified V8 agar plates using a cork borer and added to 15 ml of 2% (w/v) carrot broth amended with 15 *μ*g/ml β-sitosterol in 90 mm Petri dishes. For 1 l of 2% (w/v) carrot broth, 20 g of frozen carrots were blended for 30 s in 500 ml of ddH_2_O. The resulting mixture was then filtered through four layers of miracloth (Merck). The volume was brought up to 1 l with ddH_2_O and sterilized by autoclaving. The dishes containing agar plugs were incubated in the dark for 30 h at 22 °C. The carrot broth was then removed and the mycelial mats were washed with 15 ml of a sterile soil solution for 45 min. This had been prepared by stirring a 2% (w/v) solution of topsoil in ddH_2_O on a magnetic stirrer for 4 h at room temperature. After stirring, the solution was left to settle overnight. The solution was filtered by passage through four layers of miracloth (Merck) and then Whatman Grade 1 filter paper, before being sterilized by autoclaving. After two washes, the mycelial mats were incubated in a further 15 ml of the soil solution under continuous light for 14 h at 22 °C. The soil solution was then removed and the mycelial mats were washed with 15 ml of sterile water for 10 min. This water was removed and 15 ml of sterile water, pre-chilled to 4°C, was added. The plates were incubated at 4 °C for 20 min to stimulate zoospore release. The plates were stored at room temperature until sufficient numbers of zoospore were released (typically 1-2 h). Zoospores were pooled and counted using 2-chip disposable hemocytometers (Bulldog Bio, Portsmouth, NH, USA).

For motility and germination testing, zoospores were first diluted to a standard concentration of 5000 zoospores/ml. Next, 990 *μ*l of this suspension was added to 24-well plates containing 10 *μ*l of oxathiapiprolin at 100× the desired final concentration. For motility assays, zoospores were observed using brightfield microscopy at time intervals of 0, 10, 20, 30, 60, 90, 120, 150, 180, 210, and 240 min, or until complete loss of movement was observed. Following 240 min of observation, the 24-well plates were stored at 22°C overnight in the dark to assess germination. The following morning, zoospore germination was measured by counting 25 randomly-selected zoospores per well, with germinated spores being those with germ tubes at least twice the diameter of the spore. EC_50_ values for germination were calculated using GraphPad Prism Version 8. The total number of germinated spores was plotted against the log-transformed concentration of oxathiapiprolin.

### Oospore isolation and testing

For oospore production, three plugs (~3 mm diameter) were taken from the edge of actively growing mycelia and transferred to a 90 mm Petri dish containing 15 ml oospore growth medium (4% (w/v) carrot broth filtered through Whatman Grade 1 filter paper and supplemented with 12 *μ*g/ml β-sitosterol prior to autoclaving). The cultures were incubated in the dark for 4-6 weeks at 22 °C. The resulting mycelial mats were then transferred using sterile tweezers to 50 ml tubes and suspended in 40 ml of sterile water. To separate the oospores from mycelia, the solution was homogenised for 1 min (D160 Tissue Homogeniser, DLAB), then sonicated at 20% amplitude for 1 min (500 Watt Ultrasonic Cell Disruptor with a 5 mm probe, Sonics & Materials, Inc) and filtered through 100 *μ*m and 40 *μ*m EASYstrainers (Greiner). The filtered oospore suspension was pelleted at 1200*g* for 10 min, the supernatant removed, and the oospore pellet was resuspended in 5 ml of sterile water. Oospore numbers were estimated using disposable C-Chip haemocytometers (Bulldog Bio). Purified oospores were stored at 4 °C in the dark.

Germination assays were set up in 96-well plates with ~1000 oospores per well. A kauri root extract was used to stimulate germination. To prepare the extract, kauri roots were harvested, finely ground in a blender and added to water at 10% (w/v). This suspension was incubated overnight at room temperature with constant mixing. The resulting liquid was sterilized by passing through a 0.22 *μ*m syringe-driven filter. Germination assays contained 10 *μ*l of this kauri root extract, along with oxathiapiprolin at varying concentrations, in a total volume of 100 *μ*l. Control wells included the same concentration of DMSO as sample wells. The plates were incubated under continuous light for 5 days at 22 °C. Oospores were imaged using brightfield microscopy at 40× combined magnification.

Oospore viability assays were carried out as we have described in detail elsewhere (Fairhurst and Gerth 2021). Oxathiapiprolin at the indicated concentration was added to ~2500 oospores in a total volume of 100 *μ*l. The treated oospores were incubated at 22 °C in the dark for 2 days prior to live/dead staining. Oospores were then pelleted at 1200*g* for 10 min and the supernatant was removed. Next, 10 *μ*l of fluorescein diacetate (live stain; 200 *μ*M) was added to the oospores and incubated at 37°C in the dark for 20 h. Then 10 *μ*l of TOTO-3 iodide (dead stain; 20 *μ*M) was added and the oospores further incubated at 37°C, in the dark for 4 h. Stained oospores were imaged using a fluorescence microscope (Olympus BX63) at 40× magnification using the green filter for fluorescein (excitation 465-495 nm, emission 515-555 nm) and red filter for TOTO-3 iodide (excitation 540-580 nm, emission 590-665 nm). As a control for the live/dead staining and image analysis, oospores were rendered non-viable by heat treatment (98 °C, 24 h) before being stained and analyzed in parallel with the oxathiapiprolin-treated samples. Oospores were automatically counted by analyzing the images with CellProfiler v3.1.8 (McQuin et al. 2018) and the data were further processed and visualized as described elsewhere (Fairhurst and Gerth 2021).

### Detached leaf assays

Using a sterilized scalpel, a 2 mm incision was made 1 cm from the base of each freshly harvested kauri leaf. Three leaves were placed in a 90 mm Petri dish and each was spray coated with 200 *μ*l of oxathiapiprolin at the indicated concentration. A 6 mm diameter agar plug of *P. agathidicida* was taken from the leading edge of mycelial growth and placed over the incision on each leaf. The plates were stored at 22 °C in a 12 h light: 12 h dark cycle with a dampened paper towel to prevent leaf drying. The leaves were imaged daily for 10 days. The length of lesion spread was measured using ImageJ (https://imagej.nih.gov/ij/).

## Results

### Gene annotation and protein domain analysis

In *Phytophthora*, the biochemical target of oxathiapiprolin is the ORP1 protein (Bittner et al. 2017; Miao et al. 2018; Miao et al. 2016b; Pasteris et al. 2016). Before beginning our experimental studies, we endeavoured to identify the ORP1 ortholog in *P. agathidicida*. An unannotated, partly-assembled genome for *P. agathidicida* isolate NZFS 3772 has been reported (Studholme et al. 2016). Simple pairwise alignments identified a highly conserved ortholog of 2,973 bp contained entirely within contig 1930 of the draft genome sequence.

A previous survey identified a conserved 16 bp motif (GCTCATTYBNNNWTTY) surrounding the transcriptional start site of many oomycete genes, typically 50-100 bp upstream of the start codon (Kamoun 2003). The *P. agathidicida ORP1* gene (*PaORP1*) has a near-perfect match to this consensus (G<u>AG</u>CA<u>C</u>TCGGCCTTTC; mismatches underlined) located 153 bp upstream of the predicted start codon. In the same survey, it was also noted that introns are relatively rare in *Phytophthora* genes, but that there are conserved sequences at the 5’ and 3’ exon-intron junctions (5’-GTRNGT…YAG-3’) and a conserved intronic motif (CTAAC) important for splicing (Kamoun 2003). These features clearly define an intron sequence of 90 bp in *PaORP1*, the splicing of which also removes two in-frame termination codons. Thus, *PaORP1* encodes a 960-residue protein. The annotated sequence of *PaORP1* is shown in Supplementary Fig. S1.

The *Pa*ORP1 protein is estimated to have a molecular weight of 104.4 kDa and an isoelectric point (pI) of 6.5. Across the eukaryotes, ORP proteins are identified by a signature motif (canonically EQVSHHPP), which in turn is found within the highly-conserved OSBP-related domain (ORD) (Raychaudhuri and Prinz 2010). In the ORP1 proteins from *P. infestans, P. capsici* and *P. sojae*, the signature motif is EHTSHHPP (Andreassi et al. 2013; Miao et al. 2018). *Pa*ORP1 has the same EHTSHHPP sequence as these other *Phytophthora* orthologs and it is found at residues 728-735 (Supplementary Fig. S1). Also similar to ORP1 proteins from other *Phytophthora* species (Andreassi et al. 2013), *Pa*ORP1 is predicted to have a pleckstrin homology (PH) domain at its N-terminus, followed by a StAR-related lipid transfer (START) domain. The third domain in the primary sequence is the ORD, which is 95-97% identical to those from the other *Phytophthora* species we analyzed. The overall domain organisation of *Pa*ORP1 is shown in Fig. 1.

**Fig. 1.**
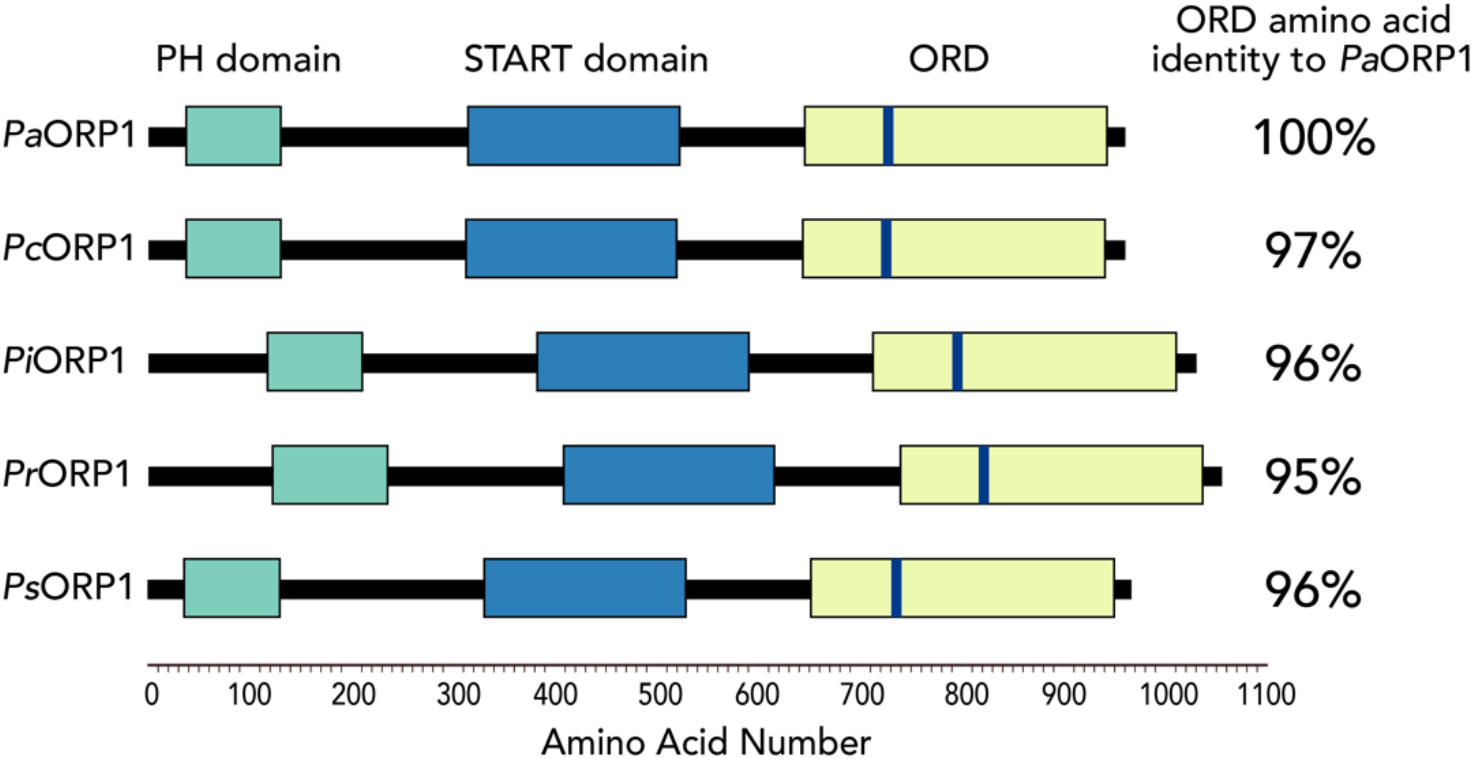
Predicted domain architecture of the *Pa*ORP1 protein and comparison with related *Phytophthora* species. *Pa*ORP1 is predicted to be a 960 amino acid protein with an N-terminal pleckstrin homology (PH) domain (green), a StAR-related lipid transfer (START) domain (blue), and the canonical C-terminal OSBP-related domain (ORD; yellow). Within the ORD is the signature motif, EHTSHHPP, indicated by a blue bar. The protein domain architecture is compared with the ORP1 proteins from *P. capsici* (*Pc*ORP1), *P. infestans* (*Pi*ORP1), *P. ramorum* (*Pr*ORP1), and *P. sojae* (*Ps*ORP1). The amino acid identity between each ORD and the *P. agathidicida* sequence is also shown.

Altogether, these bioinformatic analyses confirmed that *P. agathidicida* contains the protein target for oxathiapiprolin (*Pa*ORP1) and that, based on the high level of sequence identity to its orthologs, it was likely to be highly sensitive to this oomycide.

### Inhibitory effects on mycelia and zoospores

Oxathiapiprolin is best known for its severely inhibitory effects on mycelial growth (Gray et al. 2018; Ji and Csinos 2015; Miao et al. 2016b; Miao et al. 2016a). Its effects on mycelial growth of the two *P. agathidicida* isolates (3770 and 3772) were tested at concentrations ranging from 0.01 ng/ml to 4 ng/ml. For both isolates the EC_50_ was calculated to be approximately 0.1 ng/ml (Table 1). This indicates that *P. agathidicida* mycelia are approximately 5-to 10-fold more sensitive to oxathiapiprolin than those from well-characterized species such as *P. capsici* and *P. nicotianae* (Table 1), with a similarly low EC_50_ to that determined for *P. syringae* (Gray et al. 2018).

**Table 1.** EC_50_ values for oxathiapiprolin inhibiting selected life cycle stages of *P. agathidicida* and related species. 95% confidence intervals are listed in parentheses (*n* = 3 biological replicates).

| Pathogen | Mycelial growth EC<sub>50</sub> (ng/ml) | Zoospore germination EC<sub>50</sub> (ng/ml) |
| --- | --- | --- |
| *P. agathidicida* 3770 | 0.13 (0.11 – 0.14) | 8.5 (5.2 – 13) |
| *P. agathidicida* 3772 | 0.11 (0.096 – 0.11) | 1.8 (0.25 – 3.6) |
| *P. capsici*<sup>a</sup> | 0.68 (0.54 – 0.85) | 6.0 (0.64 – 56) |
| *P. capsici*<sup>b</sup> | 1.0 | 540 |
| *P. citrophthora*<sup>c</sup> | 0.3 | 8 |
| *P. nicotianae*<sup>c</sup> | 0.5 | NR<sup>d</sup> |
| *P. syringae*<sup>c</sup> | 0.1 | NR<sup>d</sup> |

<sup>a</sup> As reported by others (Miao et al. 2016a).
<sup>b</sup> Average across 126 isolates, as reported by others (Ji and Csinos 2015).
<sup>c</sup> Average across multiple isolates, as reported by others (Gray et al. 2018).
<sup>d</sup> NR, not reported.

The motile zoospores of *P. agathidicida* are a potential target for antimicrobials because it is they that initiate new infections by encysting and germinating on the root of the host kauri (Bellgard et al. 2016). We previously tested a range of antimicrobials and plant-derived natural products for their effects on the germination and motility of *P. agathidicida* zoospores (Lawrence et al. 2017; Lawrence et al. 2019). Oxathiapiprolin inhibits zoospore germination and (more weakly) motility in other oomycetes (Cohen 2015; Gray et al. 2018; Ji and Csinos 2015). Therefore, we set out to implement our previous methods to determine the effects of oxathiapiprolin on *P. agathidicida* zoospores.

The EC_50_ values for inhibiting zoospore germination in isolates 3770 and 3772 were 9 ng/ml and 2 ng/ml oxathiapiprolin respectively (Table 1). These values are similar to the EC_50_ reported for oxathiapiprolin acting against *P. citrophora* (Gray et al. 2018) and from a study on one isolate of *P. capsici* (Miao et al. 2016a), but approximately two orders of magnitude lower than the average determined for 126 *P. capsici* isolates in a different study (Ji and Csinos 2015).

For zoospore motility, the results were similar for isolates 3770 and 3772 across three independent preparations of zoospores (Table 2). In controls lacking oxathiapiprolin, zoospores remained motile for 180-240 min. A slight reduction in motility was first seen at 0.001 *μ*g/ml oxathiapiprolin, which was a 10-fold higher concentration than the EC_50_ for mycelial growth. Higher concentrations of oxathiapiprolin reduced zoospore motility to a greater degree, but even at 1 *μ*g/ml (10,000-fold higher than the mycelial EC_50_) zoospores remained motile for 60 min (Table 2). Comparable results have been observed for *Pseudoperonospora cubensis*, the oomycete pathogen of cucurbits. At 30 min after treatment with 5 *μ*g/ml oxathiapiprolin, 25% of *P. cubensis* zoospores remained motile (Cohen 2015). On the other hand, we previously showed that copper (II) fungicides (Lawrence et al. 2017) and New Zealand native plant-derived flavonoids (Lawrence et al. 2019) rapidly immobilized *P. agathidicida* zoospores at concentrations that were significantly lower than their EC_50_ values for mycelial growth inhibition.

**Table 2.** Oxathiapiprolin inhibition of zoospore motility. The mean time until complete loss of motility of all zoospores is listed for each oxathiapiprolin concentration (*n* = 3 biological replicates).

| [Oxathiapirolin], $\mu\text{g/ml}^a$ | Isolate 3770 | Isolate 3772 |
| --- | --- | --- |
| 0 | 180 min | 240 min |
| 0.001 | 150 min | 210 min |
| 0.01 | 90 min | 120 min |
| 0.02 | 90 min | 90 min |
| 0.04 | 90 min | 90 min |
| 0.08 | 60 min | 90 min |
| 0.16 | 60 min | 60 min |
| 1.0 | 60 min | 60 min |

<sup>a</sup> Oxathiapirolin was dissolved in DMSO; therefore DMSO was added to all wells at a final concentration of 1% (v/v) to control for any solvent effects.

### Inhibitory effects on oospores

Oospores are key survival propagules in soil for many *Phytophthora* species (Erwin and Ribeiro 1996), particularly those such as *P. agathidicida* that do not produce chlamydospores (Weir et al. 2015). They can lie dormant for years and germinate under the right conditions, leading to the formation of sporangia, release of zoospores, and thus new infections. The dormant oospores are easy to transfer between sites, such as on shoes bearing contaminated soil, making them an important consideration for minimizing disease spread. Oxathiapiprolin is highly effective at preventing mycelial mats of *P. nicotianae* from producing oospores (Gray et al. 2018). However, it is technically challenging to produce pure preparations of oospores and therefore, to test the effects of oomycides upon them. To the best of our knowledge, no-one has investigated the direct effects of oxathiapiprolin on oospores themselves. Here, we have used our recently developed protocols for producing purified *P. agathidicida* oospores, assessing viability, and for triggering their germination *in vitro*, to test the effects of oxathiapiprolin on this key lifecycle stage.

First, we assessed the effect of oxathiapiprolin on oospore germination. Oxathiapiprolin concentrations from 0.1 *μ*g/ml to 10 *μ*g/ml completely inhibited germination (Fig. 2A). On the other hand, live/dead staining showed that oxathiapiprolin did not render oospores non-viable when applied at the same concentrations (Fig. 2B). That is, *P. agathidicida* oospores can withstand oxathiapiprolin treatment but cannot successfully germinate in its presence. Oospores of both *P. agathidicida* isolates (3770 and 3772) behaved identically in these tests.

**Fig. 2.**
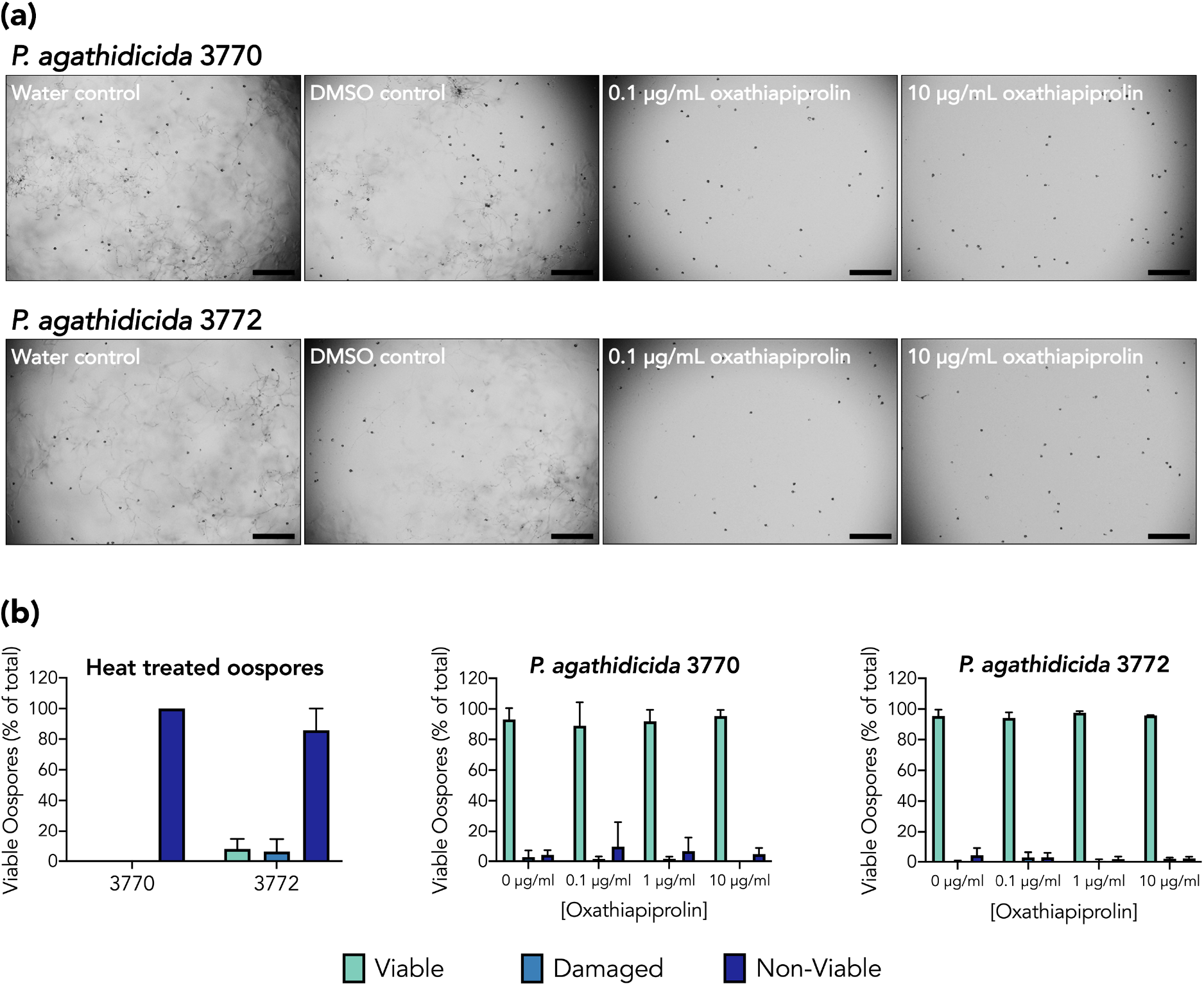
Effects of oxathiapiprolin on *P. agathidicida* oospores. (**a**) Inverted light microscopy images of oospores triggered to germinate by the addition of kauri root extract, in the presence or absence of oxathiapiprolin. New mycelial mats from germinated oospores could not be captured in a single focal plane so they appear as a combination of fibrils and hazy shadows in the water-only and DMSO controls. Scale bar = 250 *μ*m. (**b**) Results of live/dead staining for oospore viability. The left panel shows the results from heat treating oospores of each *P. agathidicida* isolate, to verify the viability screening method. The other panels show the effects of treating each isolate with varying concentrations oxathiapiprolin. Error bars show the standard deviation from *n* = 3 independent oospore preparations.

### Detached Leaf Assays

Detached leaf assays provide a simple measure of infectivity on host plants (Goth and Keane 1997; Pettitt et al. 2011). They can be used to assess whether test compounds are preventative and/or curative (Cohen et al. 2018). Here, we tested oxathiapiprolin for its preventative effects. Kauri leaves were damaged and then sprayed with oxathiapiprolin at different concentrations, before being inoculated with each isolate of *P. agathidicida*. The onset of infection for untreated kauri leaves was observed at 4-5 days for both isolates. In contrast, pre-treatment with low concentrations of oxathiapiprolin (0.1 *μ*g/ml and 1 *μ*g/ml) delayed the onset of infection until day 8 and day 9, respectively (Fig. 3). The highest concentration of oxathiapiprolin (10 *μ*g/ml) completely inhibited infection during the ten-day observation period.

**Fig. 3.**
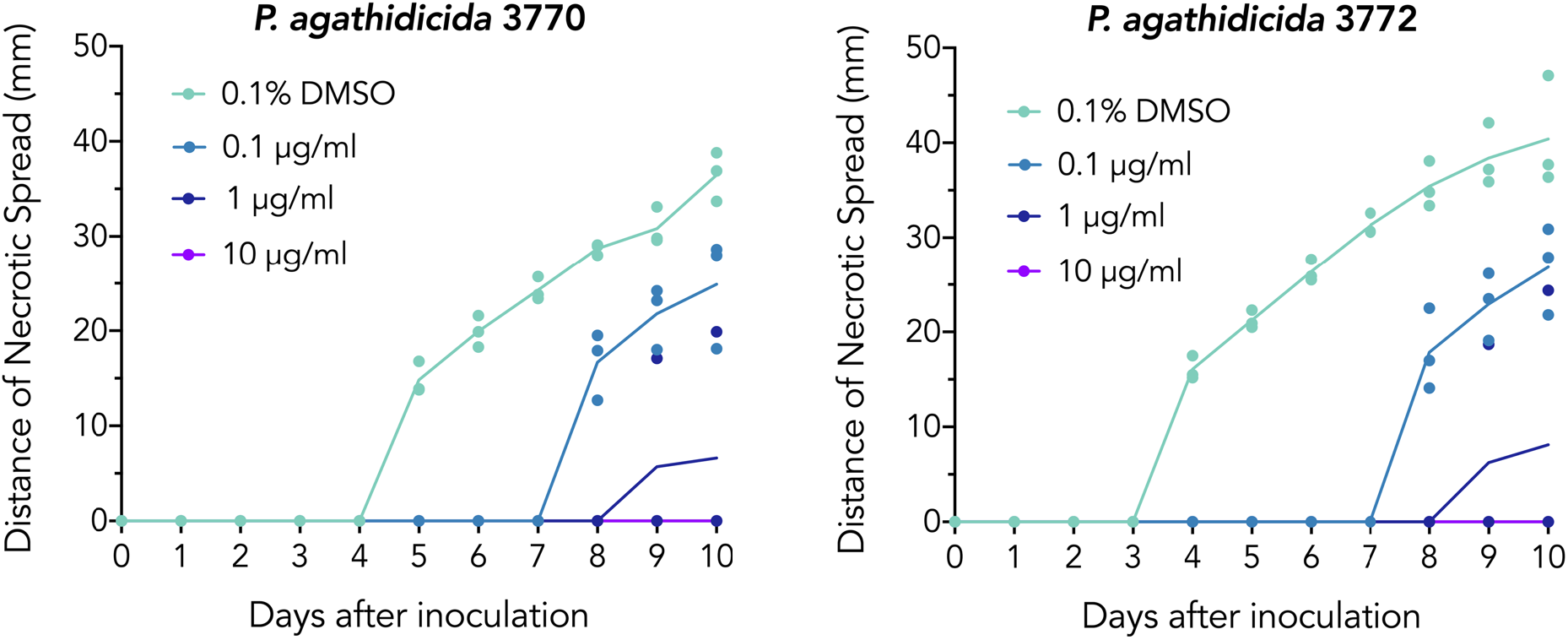
Preventative effects of oxathiapiprolin on infection of kauri leaves. Detached leaf assay results for *P. agathidicida* isolates 3770 (left panel) and 3772 (right panel). The oxathiapiprolin concentrations used to pre-treat leaves are indicated and each data point is shown for *n* = 3 independent replicates at each concentration. The connecting lines indicate means of the necrotic spread observed for each treatment. Leaf necrosis indicates *P. agathidicida* infection; therefore, reduced or absent necrosis indicates that the treatment has a preventative effect.

## Discussion

Oxathiapiprolin is a novel oomycide that is highly active against many species of *Phytophthora* at extremely low concentrations (Belisle et al. 2019; Gray et al. 2018; Ji and Csinos 2015; Pasteris et al. 2016). In this study we sought to determine its effectiveness against multiple lifecycle stages of *P. agathidicida*, the causative agent of kauri dieback disease. Similar to observations for other *Phytophthora* species, oxathiapiprolin was effective against mycelial growth and zoospore germination at concentrations far below 1 *μ*g/ml (Table 1). Indeed, the measured EC_50_ values for mycelial growth indicate that *P. agathidicida* mycelia are even more sensitive to oxathiapiprolin than well-studied species such as *P. capsici* and *P. cinnamomi* (Belisle et al. 2019; Ji and Csinos 2015; Miao et al. 2016b). Oxathiapiprolin was also effective at delaying or preventing the infection of kauri leaves, albeit at higher concentrations (Fig. 3). On the other hand, it failed to render zoospores immotile in less than an hour (Table 2). Overall, its inhibitory effects were comparable for the two geographically distinct *P. agathidicida* isolates 3770 and 3772.

In assessing the effects of oxathiapiprolin on the *P. agathidicida* lifecycle, a particular focus was how it affects oospores. This is an often-overlooked aspect of oomycide efficacy. If a compound is effective at inhibiting mycelial growth *in planta*, yet ineffective at inhibiting oospore germination and/or viability, disease can readily re-emerge from tissues containing oospores, or new infections can rapidly initiate from oospore-containing soil. The only previous study of *P. agathidicida* oospores showed that viability is largely unaffected by Trigene (a commercial disinfectant containing halogenated tertiary amines), salt water immersion or various pH treatments, with substantial heat treatment being the only effective measure for reducing viability *in vitro* (Dick and Kimberly 2013). Here we found oxathiapiprolin to be highly effective at inhibiting germination, but like these other treatments it was ineffective at reducing oospore viability (Fig. 2). Our results highlight the importance of assessing as many lifecycle stages as possible when testing new oomycides, in order to obtain a complete picture of their effectiveness.

Trunk injection of phosphite is the only chemical treatment currently being used against confirmed cases of kauri dieback (Bradshaw et al. 2020; Horner et al. 2015). Here we have conducted the first tests to establish whether oxathiapiprolin should also be considered. Currently in New Zealand, oxathiapiprolin (sold as Zorvec Enicade: https://www.corteva.co.nz/products-and-solutions/crop-protection/zorvec-enicade.html) is only approved for the control of *Peronospora destructor* (downy mildew) in bulb onion. However, our results show that oxathiapiprolin is also highly effective against several key lifecycle stages of *P. agathidicida*. By acting at sub-microgram per millilitre concentrations to inhibit mycelial growth, zoospore germination and oospore germination (albeit *in vitro*), oxathiapiprolin emerges as a promising candidate for further testing. Inhibiting zoospore and oospore germination highlights its potential as a preventative agent, whereas its extreme effectiveness against mycelial growth suggests it may also act curatively, even in large trees. Future studies should focus on determining the optimal dosage, application method, uptake and overall efficacy *in planta*. Ultimately, field trials of oxathiapiprolin against confirmed cases of kauri dieback will be a critical part of the ongoing efforts to protect kauri, which are unique taonga (treasures) of New Zealand.

