## Supplementary Figure 1 for "Assessing the effectiveness of oxathiapiprolin towards *Phytophthora agathidicida*, the causal agent of kauri dieback disease"

```
GAGCACTCGGCCTTTCATCCCCTCGCGCGTACGTGTCGAAACCTCTTGGTACTCGAGAGCCTCGACAGGGAGCAGGGGCCGCAGACCTTAGCACACCACGAAGGACCAAGGCAACGGCCA 120
                                                M  Q  A  L  Q  D  A  Q  Q  R  F  N  D  L  V  N  N  D
CAGCGCCAACGGCGGCGACGCGGCTCGCGAACGCAAGCGGGACCGAGCCATGCAGGCGCTTCAAGACGCGCAGCAGCGCTTTAATGACCTGGTTAACAATGACGTACGTTAACTTGTTAT 240
                                                                                 W  P  E  R  V  P  V  E  S  M  P  D  Y  D  P  S
CCCCTGTTATCAAAAAAAGATAAACCCTAACATGACAATTGACAAACCCTAACTTTATATTTCTTCTTTGCAGTGGCCTGAGCGGGTCCCCGTGGAGTCCATGCCGGACTACGACCCGAG 360
 Y  M  K  E  G  F  L  Q  K  K  G  Q  R  L  K  G  W  K  R  R  W  F  V  C  D  G  R  T  L  S  Y  Y  I  S  R  K  D  R  K  P
CTACATGAAGGAAGGGTTCTTGCAAAAGAAGGGCCAGCGTCTTAAAGGCTGGAAGCGCCGCTGGTTCGTCTGTGACGGCAGGACGCTGTCCTACTACATTTCCAGGAAAGACCGCAAACC 480
 N  A  V  I  P  L  E  G  C  T  V  Q  D  G  G  L  S  E  T  W  N  S  P  R  I  Y  L  T  D  P  A  T  G  I  M  Y  C  L  S  A
CAATGCGGTGATCCCATTAGAAGGCTGCACAGTGCAGGACGGAGGGCTCAGTGAGACGTGGAACTCCCCTCGCATTTATTTGACGGACCCAGCCACCGGTATCATGTACTGCTTGTCGGC 600
 E  E  G  V  V  V  T  Q  W  L  D  V  L  R  V  A  V  A  R  V  N  N  G  S  A  A  S  E  T  N  A  N  S  S  S  S  A  P  S  S
TGAAGAAGGGGTCGTGGTGACTCAGTGGCTGGATGTGCTCAGGGTGGCGGTGGCTAGAGTGAACAATGGGTCAGCTGCTAGTGAGACCAATGCCAATTCCAGCAGTTCTGCACCGTCGTC 720
 Q  G  S  S  R  R  P  Q  R  T  Q  A  Q  R  L  P  S  S  S  D  D  E  D  S  R  A  H  L  K  R  A  A  S  L  G  G  P  A  Q  P
ACAGGGATCAAGTAGAAGACCACAGAGAACGCAGGCTCAGAGGCTCCCGTCGTCTTCGGATGACGAAGACAGTCGAGCACATTTGAAGCGTGCGGCGTCGCTGGGAGGACCCGCGCAGCC 840
 R  T  T  T  L  K  S  A  S  S  A  V  T  S  S  N  T  V  V  N  G  E  G  K  R  A  T  R  M  T  S  A  P  S  A  V  S  T  S  N
TCGTACGACGACGTTGAAGTCGGCGTCGTCGGCGGTGACGAGTTCGAATACTGTTGTGAATGGAGAGGGCAAGCGTGCGACGAGGATGACGTCGGCACCGTCGGCGGTGAGTACTTCGAA 960
 S  S  A  V  Q  P  A  Q  H  H  H  H  R  V  H  R  T  K  T  Q  R  L  P  T  T  I  A  L  E  N  E  L  S  H  G  L  D  V  L  E
CAGCAGTGCAGTGCAGCCGGCGCAGCATCATCATCATCGAGTTCATCGTACAAAGACGCAGCGACTGCCGACGACGATTGCGCTGGAGAATGAGCTTTCGCATGGGTTAGATGTGCTAGA 1080
 A  L  L  C  G  H  S  A  T  G  S  S  S  S  I  R  N  H  V  V  F  R  P  I  G  A  L  N  G  V  L  R  S  I  G  T  D  S  S  S
GGCACTGCTGTGCGGTCATAGCGCGACAGGATCGTCGTCTTCGATCCGCAATCACGTTGTGTTCCGACCGATTGGCGCCTTGAACGGTGTGCTACGCAGCATTGGAACGGATTCAAGCTC 1200
 G  K  Q  Y  A  R  A  S  V  V  L  P  V  S  S  E  V  A  A  I  L  L  A  D  H  G  R  R  A  E  W  D  V  H  F  P  Q  S  S  H
TGGCAAGCAATATGCGCGAGCGAGTGTTGTGTTGCCCGTGTCTTCTGAAGTTGCGGCGATTTTGCTGGCGGATCACGGGCGCCGTGCTGAATGGGATGTACACTTCCCACAGTCATCGCA 1320
 V  A  N  F  D  D  A  T  D  L  V  H  L  S  S  G  S  F  A  Q  V  Q  C  T  K  P  F  V  A  P  H  V  A  A  A  A  C  A  L  C
CGTAGCCAACTTTGACGACGCGACGGACCTCGTGCATTTGTCGAGTGGAAGCTTCGCACAGGTCCAGTGCACTAAACCGTTTGTGGCTCCTCATGTGGCTGCAGCGGCGTGCGCCTTGTG 1440
 A  A  L  F  S  G  A  S  S  W  E  G  L  L  T  A  M  V  Y  A  A  A  I  G  G  I  V  S  S  I  D  Y  S  S  L  T  A  P  R  D
TGCGGCACTTTTCTCGGGCGCTTCATCGTGGGAAGGATTGCTGACCGCTATGGTGTACGCTGCTGCCATTGGCGGCATTGTGAGCAGCATCGATTACAGTTCATTGACGGCGCCCCGTGA 1560
 L  V  I  L  R  H  V  R  E  S  A  T  P  D  S  Q  D  S  G  D  D  K  S  V  D  E  M  G  Q  S  V  A  L  I  L  E  K  S  V  V
CCTAGTGATTCTTCGCCACGTGCGTGAGTCTGCTACACCCGATTCACAGGATTCAGGCGACGACAAGTCTGTTGATGAGATGGGCCAGTCTGTGGCGCTCATCCTGGAGAAGTCTGTGGT 1680
 N  E  L  K  P  V  I  S  G  T  V  R  A  H  V  G  L  S  G  W  L  L  E  P  V  D  S  G  R  A  T  L  A  T  Y  I  T  D  L  D
CAACGAGCTGAAGCCGGTCATCTCGGGCACTGTGCGTGCACATGTAGGACTGAGCGGCTGGTTGCTGGAACCGGTGGATTCAGGCCGCGCTACGCTGGCCACTTACATTACGGATTTGGA 1800
 M  K  G  W  L  S  P  S  T  R  Q  R  F  L  L  S  R  L  D  C  V  S  V  L  S  E  Y  V  N  Q  A  Q  L  C  G  S  E  L  G  F
TATGAAGGGGTGGTTGTCGCCCTCCACGCGCCAAAGATTCTTGCTGTCGCGCCTGGACTGCGTGTCTGTGCTGAGTGAATACGTTAACCAAGCCCAGCTGTGCGGATCCGAGCTTGGCTT 1920
 G  G  G  L  D  E  D  G  D  G  E  Y  E  T  R  S  V  G  Y  V  D  T  S  E  G  S  A  L  G  D  V  V  D  G  E  S  P  E  I  F
TGGTGGTGGCTTGGACGAAGATGGTGACGGGGAGTACGAAACTCGTAGCGTAGGTTACGTAGACACTAGCGAAGGGAGCGCTCTTGGAGATGTTGTGGATGGCGAATCCCCCGAGATTTT 2040
 H  P  K  T  Y  M  R  G  M  M  P  L  P  S  G  G  L  K  L  I  D  K  E  V  A  K  K  Q  G  G  V  V  K  D  V  I  K  S  A  G
CCACCCCAAGACATACATGCGTGGCATGATGCCGTTGCCTAGCGGTGGCTTGAAATTGATCGACAAGGAAGTCGCCAAGAAGCAGGGTGGCGTGGTGAAGGATGTCATCAAATCGGCAGG 2160
 A  K  I  L  E  G  K  S  A  V  S  L  S  L  P  V  R  I  F  E  P  R  T  N  L  E  R  V  C  D  L  M  L  Y  A  P  T  F  L  N
AGCAAAAATCCTGGAGGGCAAGTCGGCTGTGAGTCTATCGCTGCCTGTGCGTATTTTCGAGCCTCGCACCAATCTGGAACGTGTGTGCGATTTAATGCTGTACGCGCCGACGTTCCTGAA 2280
 V  A  H  A  Q  N  D  A  L  E  R  F  K  Y  V  M  T  F  A  V  A  G  L  H  H  S  I  G  Q  L  K  P  F  N  P  I  L  G  E  T
CGTTGCACACGCGCAGAACGATGCTTTAGAGCGATTCAAGTACGTCATGACATTTGCCGTGGCTGGTCTGCACCACAGTATTGGACAGTTGAAGCCGTTTAACCCCATCTTGGGTGAGAC 2400
 F  Q  S  T  L  N  D  G  T  D  V  S  C  E  H  T  S  H  H  P  P  I  S  N  F  Q  F  T  G  E  K  Y  S  I  A  G  F  V  L  W
GTTCCAGTCTACGCTGAACGATGGTACAGATGTAAGTTGCGAACACACGAGCCATCACCCGCCTATCAGTAACTTCCAGTTCACCGGAGAAAAGTACTCCATCGCTGGTTTTGTGCTGTG 2520
 H  A  S  M  S  V  K  S  N  A  M  L  N  T  N  K  G  P  V  R  V  T  F  P  D  S  E  G  L  P  G  T  T  I  E  Y  N  L  P  Y
GCATGCGAGCATGAGCGTGAAGTCGAATGCAATGCTCAACACAAACAAGGGGCCTGTGCGCGTGACGTTCCCCGACTCTGAAGGCCTTCCTGGAACGACCATCGAGTACAACCTACCGTA 2640
 L  Q  I  G  G  L  L  W  G  D  R  T  V  D  I  M  G  N  M  V  F  E  D  K  K  N  H  L  Q  C  E  L  R  L  N  P  D  A  K  S
CTTGCAGATTGGTGGGTTGCTTTGGGGAGACCGTACCGTTGACATCATGGGGAATATGGTGTTTGAAGACAAGAAGAACCATCTACAGTGCGAACTGCGCCTCAATCCGGATGCCAAGTC 2760
 G  M  G  G  M  F  S  S  S  K  T  P  T  D  S  L  R  G  V  I  L  D  T  S  V  S  P  P  R  E  I  C  D  V  S  G  S  W  L  H
AGGAATGGGCGGAATGTTTTCTAGCTCTAAAACCCCGACGGATTCGTTGCGTGGTGTGATCTTGGACACTTCTGTGTCTCCACCGCGTGAGATTTGTGACGTCTCGGGCTCCTGGCTGCA 2880
 D  L  V  F  G  N  K  T  Y  W  S  I  N  K  H  H  S  G  Y  M  V  P  Y  P  E  N  K  I  L  A  S  D  S  R  Y  R  E  D  L  H
TGACCTCGTGTTCGGCAACAAGACGTACTGGAGCATCAACAAACACCACAGCGGCTACATGGTACCGTACCCCGAGAACAAGATTTTGGCGTCCGACTCTCGGTACCGCGAAGACCTGCA 3000
 Y  L  A  A  G  D  L  D  E  S  Q  E  W  K  V  K  L  E  V  L  Q  R  A  D  R  K  A  R  L  D  G  R  R  P  N  H  W  S  F  R
CTATCTGGCAGCAGGCGACCTGGATGAATCGCAAGAGTGGAAGGTAAAGCTGGAGGTTCTGCAGCGCGCTGATCGCAAGGCGCGTCTGGACGGCCGACGTCCCAACCACTGGTCTTTCCG 3120
 S  S  A  G  G  H  *
GAGTTCTGCTGGTGGTCACTAA 3142
```

**Supplementary Figure S1. Annotated *PaORP1* gene sequence.** Red indicates the consensus sequence of a predicted transcriptional start site; black is a 5' non-coding region; green is exon 1; grey is intron 1 with bold underline indicating the conserved intronic motif; blue is exon 2 with the bold and underlined region indicating the conserved ORD sequence. The signature motif of the ORD is highlighted in yellow. The amino acid sequences for the two exons are indicated in bold above each exon.
